# Modulation of articulation muscles during listening and reading: a matter of intention to speak out loud

**DOI:** 10.1101/728485

**Authors:** Naama Zur, Zohar Eviatar, Avi Karni

## Abstract

The articulation of speech sounds is often contingent on the intention to subsequently produce other sounds (co-articulation). Thus, intended acts affect the way current acts are executed. We show that the intention to subsequently repeat a short sentence, overtly or covertly, significantly modulated articulatory muscle activity already during speech perception or reading (input interval) and when delaying response (i.e., prior to production). Young adults were instructed to read (whole sentences or word-by-word) or listen to recordings of sentences to be repeated afterwards, either covertly or overtly. Surface electromyography (sEMG) recordings showed different patterns of articulatory muscle modulation in the two articulatory muscles measured – orbicularis oris inferior (OOI) and sternohyoid (STH). In the OOI, activity increased relative to baseline during speech perception or reading in both intended output conditions (overt and covert); in the STH, articulatory muscle activity decreased, during the input intervals, in both intended output conditions. However, the modulations in EMG activity were contingent on the intention to subsequently repeat the input overtly or covertly, so that activation in the OOI, and inhibition in the STH, were significantly more pronounced when overt responses were intended. Input modality was also a factor; immediately before overt responses, activity in the OOI muscle increased for listening and word-by-word reading, but not in reading whole sentences. The current results suggest that speech perception and articulation interact already during the input phase, listening or reading, reflecting the intended responses. However, this interaction may differentially affect facial-articulatory and laryngeal control mechanisms of speech production.

## 1. Introduction

Whether and how motor representations of speech are integral to speech perception has been debated for many decades. One perspective posits that the motor system plays a functional role both in speech production and in speech perception (Liberman, 1957; Liberman, 1967; Wilson et al., 2004; Pulvermüller and Fadiga, 2010; Yuen et al., 2010; Skipper et al. 2017)^1–6^. A related position assumes an interaction between the perception of speech and the motor articulatory system, although the actual activation of the motor system is not necessary for perception (Hickok & Poeppel, 2007, Poeppel & Hickok, 2004; Scott et al., 2009)^7–9^. For example, according to feedback control models of speech production, there are pathways both for activation of motor speech systems from sensory input (forward prediction pathway) and for mutual activation of auditory speech systems from motor activation (feedback correction pathway) (Hickok et al., 2011; Guenther & Hickok, 2016)^10,11^. Thus, the state of articulatory musculature may be potentially affected during speech perception (Galantucci et al., 2006; Glenberg & Gallese, 2012; Hickok et al., 2011)^12–13,10^. Current notions of whether and how motor representations of speech contribute to speech perception do not differentiate between the effects of different language input modalities. The primary modality for language is auditory, but in literate individuals the interpretation of visual written language also becomes automatic (Logan, 1997; Augustinova & Ferrand, 2014) ^14–15^. Input-modality specific effects on articulatory muscles can lend support for the possibility of more than a single mode of interaction between language perception and speech production. In the present study, the state of articulatory muscles during speech perception – language input phase - was recorded to test the conjecture that the articulatory musculature is affected by the intention to subsequently produce covert vs. overt speech but also by the mode of language input (listening, reading). To this end, a repetition task was developed, requiring the delayed production of either covert or overt speech in response to sentences presented as auditory or visual input, and the modulation of articulatory musculature activity preceding sentence repetition was examined.

We recorded surface electromyography (sEMG) over the Orbicularis Oris Inferior (OOI) and the Sternohyoid (STH) muscles throughout the repetition task, as shown in panel A of Figure 1. The motivation for including the STH stemmed from the crucial role that laryngeal muscle activity and activity in the Laryngeal Motor Cortex (LMC) have in learned vocal behaviors specifically speech production and singing (Jürgens, 2002; Simonyan, Ostuni, Ludlow, and Horwitz, 2009) ^16, 17^. Symonian (2014)^18^ suggests that LMC topography and connectivity in humans is potentially the underlying cause for the unique human ability to produce highly controlled motor outputs characteristic of speech.

**Figure 1:**
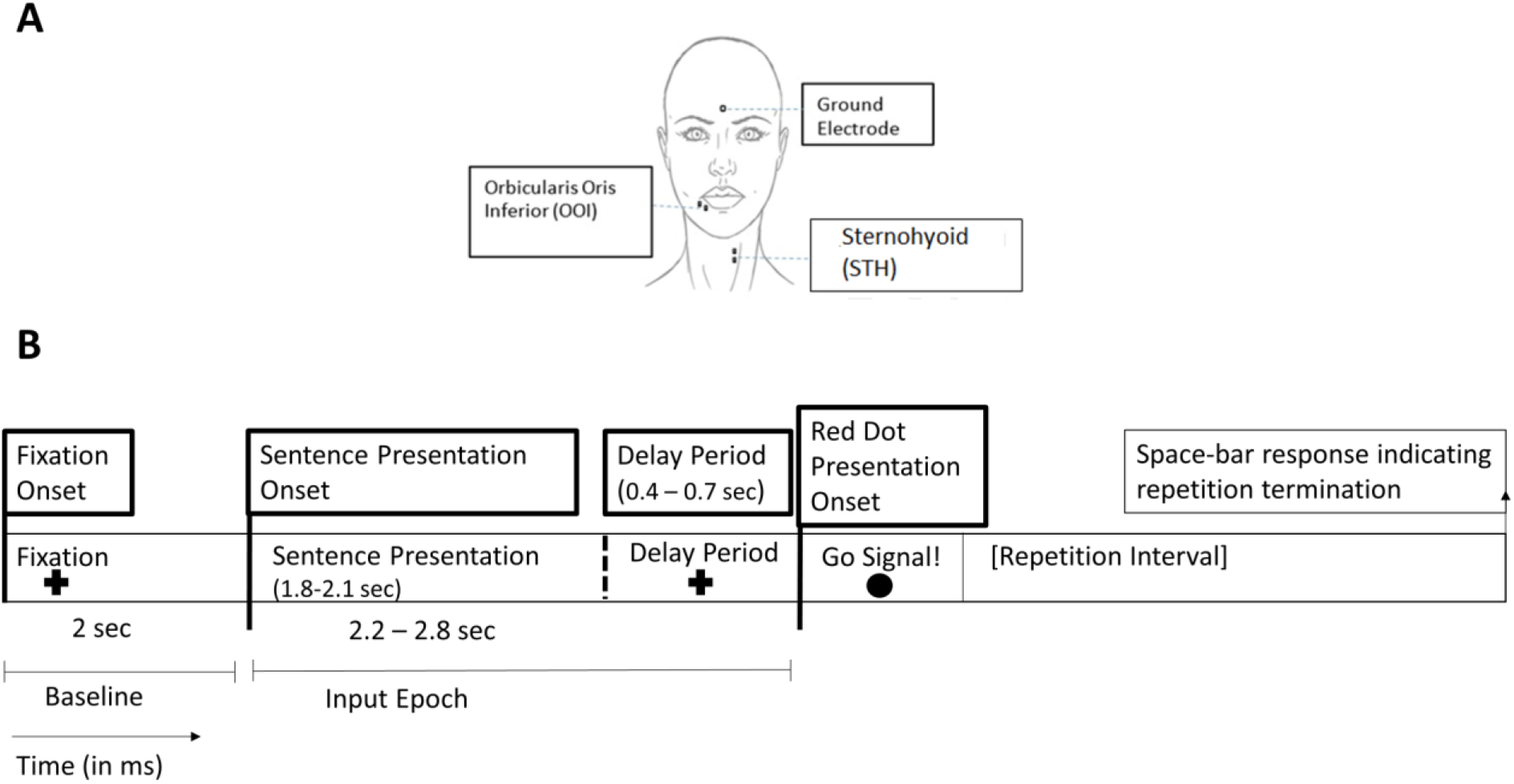
Placement of the sEMG electrodes and the timeline of each trial. **Panel A**: sEMG was recorded from the Orbicularis Oris Inferior (OOI) and Sternohyoid (STH) muscles with 4-mm miniature Beckman Ag/AgCl electrode pairs (gel filled). Skin sites were prepared with Nuprep (Weaver and Company, Aurora, CO) before electrode application. **Panel B**: The sequence of events on each trial – a fixation target (2 sec) preceded stimulus presentation (fixation segment); a stimulus presentation segment of variable duration (with randomly chosen time jitter wherein the fixation cross reappeared, total 2.2 – 2.8 sec); a Go signal (0.5 sec) and a response interval (ending with a space bar press).

Previous studies have shown subtle activity in speech musculature using sEMG during verbal mental imagery, silent reading, and silent recitation (Jacobson, 1931; Livesay et al., 1996; McGuigan, & Dollins, 1989)^19–21^. Also, activity in the OOI and the STH muscles was significantly higher when patients with schizophrenia reported hallucinations (Rapin et al., 2013)^22^. However, a ‘premotor silent period’, that is, a brief inhibition relative to baseline has been found before the initiation of both real and imagined movements in tasks unrelated to language (Conrad et al., 1983; Richartz et al., 2010; Kolářová et al., 2016) ^23–25^. Thus, the approach to data analysis in the current study was to compare the direction (excitation or inhibition) and magnitude of the modulation of activation in articulatory muscles in the different input and intended response conditions, during the input, i.e., perceptual epoch; before sentence repetition was cued (panel B of Figure 1).

## 2. Materials and Methods

### 2.1 Participants

Twenty undergraduate students (5 males) 21-35 years old, participated in the study. All were healthy native speakers of Hebrew, right handed, and had normal or corrected-to-normal vision and normal hearing. Informed consent forms were obtained from every participant prior to their participation. The study was approved by the Ethical Review Committee of the University of Haifa, and all methods were carried out in accordance with relevant guidelines and regulations, complied with the standards defined in the Declaration of Helsinki. The participants received either course credit or monetary compensation for their time (70 NIS).

### 2.2 Stimuli

Given that stimulus recognition in the auditory and visual modality can be temporally distinct (Whiting et al., 2014)^26^, there were two visual presentation conditions: a visual whole sentence presentation condition (the whole sentence was presented onscreen), and a visual word-by-word condition in which each word of the sentence was presented separately (at the center of the screen). The visual word-by-word condition was to test whether a manipulation within the visual modality, where the linguistic input is presented incrementally in time (as in auditory presentation) would differ, in terms of the motor activity modulation of articulatory musculature, from the visual whole sentence presentation condition.

The experiment included a set of 60 sentences (3 words, subject-verb-object). There were 3 sets of 20 sentences (each set containing 96 syllables of 100 possible syllables in Hebrew). The stimulus sets were counterbalanced across participants and the three input mode conditions (auditory, visual whole sentence, visual single word sentence). Visual sentence presentation was in Ariel font (size 20), black - on a gray background.

The duration of visually presented sentences was equated to the duration of the corresponding recorded sentences (Audacity(R) Version 2.0.0. (Audacity Team, 2012)); for word-by-word visual stimuli, each word was presented separately, at the center of the screen, for a duration proportionate to the duration of its voiced production.

### 2.3 Apparatus

Surface EMG data were recorded using passive electrodes (Impedance was ≥30Ω) (See Figure 1) at a sampling rate of 1000 Hz and amplified with a multichannel BioNex 8-slot chassis (Mindware Technologies, Grahanna, OH) equipped with a two BioNex 4-channel bio potential amplifier (Model 50-371102-00) (Figure 1). Data were viewed using MindWare acquisition software BioLab 2.4.

The experiment was designed using E-Prime 2 professional software. and run on a HP PC with manual responses recorded using a keyboard and vocal responses through a Lavlier microphone (Alto Professional wireless microphone system) and an e-prime compatible response box. A Dell UltraSharp U2412M color monitor and headphones (Sony Stereo-Headphones, MDR-XD100) were used for visual and auditory sentence presentation, respectively. Participants were video recorded continuously.

### 2.4 Experimental Design

The experimental design was a 2 (Intended Output: Covert and Overt) X 3 (Input Type: Auditory (A), Visual Whole Sentence (V1), and Visual Word-by-Word (V2)) within subject comparison design. Input conditions (A, V1, V2) were mixed and counterbalanced within each block. Each target sentence was presented twice (for overt and for covert repetition) but never in the same input modality. There were separate blocks wherein either a covert or an overt repetition was required. The covert block always preceded the overt blocks (panel B of Figure 1).

### 2.5 Procedure

Before the presentation of each sentence, a fixation cross appeared at the center of the screen for 2 seconds. On each trial, participants were presented with the sentence to be repeated (in either the visual or auditory modality) but were instructed to repeat the target sentence only after given a visual ‘go’ cue (Figure 1). Thus, each trial was divided into a baseline fixation epoch, an ‘input’ epoch, which was the sentence presentation part of the trial, and an ‘output’ epoch, after the cue to repeat. This analysis reduces the need for a temporally offset, separate, baseline condition such as maximal volitional contraction (MVC) condition to control for these large differences (Stepp 2012)^27^, because for each participant, the individual’s specific baseline for the ongoing trial was used. Thus, we compared musculature activity at a given interval in each trial to an immediately preceding, non-verbal baseline. The difference measure enabled us to conduct a within-subject analysis decreasing the variance due to individual differences, bypassing the ubiquitous common methodological problem with sEMG of individual differences in the raw signal.

The sentence to be repeated followed in one of three modes: 1) auditory, via headphones (the fixation cross stayed on screen), 2) visual, sentence in full at the center of the screen (no audition), 3) visual, word by word presentation of the sentence at the center of the screen (no audition). In the first block, participants were instructed that repetition was to be covert (silent repetition); in the second block repetition was to be overt (repetition out loud). Immediately following sentence presentation offset, a fixation cross was presented at the center of the screen for a duration of 400-700 ms (jitter), until the onset of a red dot. The red dot cued response initiation (Go cue) and was presented at the center of the screen until the participant pressed the “space” key (end of response).

Response times (RT) were measured from the onset of the red dot to “space” key press. Six trials with responses over 5 seconds or shorter than 200 ms were excluded. Inaccuracy of production (overt repetition block) occurred in eight trials (out of 1200 responses); these trails were excluded.

### 2.6 EMG data processing

Preprocessing was done using MATLAB R2018b (MathWorks Inc.). The sEMG raw data were rectified by absolute value and fed into a 20–450 Hz Butterworth band-pass filter (Butter, filtfilt, MATLAB). Additional movement artifact removal was done by offline inspection of the video recording. Swallowing was determined using both the video recordings (for physical movement of STH electrodes) and irregular peaks in the signal.

About 4.7% of the trials (113/2400 trials) were excluded from analysis due to technical and movement artifacts.

In the analysis, we first addressed the dynamics of muscle activity during the fixation interval, by comparing the RMS values of two intervals of 200 ms each, one from the beginning of the fixation epoch (Fix1) and one from the end (Fix2) (Figure 3). The goal of this analysis was to confirm that the difference measure of activity modulation between intended output conditions (covert and overt) was comparable. Therefore, we tested whether there would be a modulation throughout the fixation epoch as a function of the two intended output conditions (covert vs. overt). In both conditions and in both muscles there was lower activation levels in Fix2 compared to Fix1; in the OOI, the difference between the two fixation intervals was greater in the Overt than in the Covert condition. We next compared muscle activity (relative to Fix2) in four intervals of 200 ms each (Figure 5) within the stimulus presentation segments: an interval starting at the onset of sentence presentation (interval A), an interval at the end of sentence presentation – from 300 ms to 100 ms prior to the end of the presentation (interval B), an interval starting at the offset of sentence presentation (interval C), and an interval just before the repetition cue (interval D). RMS values of sEMG activity (μV) were computed for each of the intervals, and a relative activity measure was computed by subtracting the RMS (μV) value of Fix2 from each of the RMS(μV) values of the intervals.

### 2.7 Statistics

Three-way within-subject ANOVAs were conducted the all three within-subject factors in the design (Input Type, Intended Output, and Interval), and multiple comparisons were controlled using the Bonferroni correction. Mauchly's Test of Sphericity was used to check for violations in sphericity in each factor. Violations of sphericity were corrected either with the Greenhouse-Geisser or Huynh-Feldt corrections. One-sample t-tests were run on all conditions of the design to examine whether they were significantly different from baseline. Statistical analyses were computed with IBM SPSS Statistics for Windows, version 23 (IBM Corp., Armonk, N.Y., USA).

## 3. Results

The panels in Figure 2 present examples of the raw sEMG recordings (in μV) from both the OOI and STH muscles recorded from one of the participants (a representative trial from each of the 6 study conditions). For each trial, we computed the average values for the root mean square (RMS) of the sEMG activity (μV). The values were extracted from different segments (epochs) of each trial: 1) fixation epoch – the last 200 ms of the 2 second fixation cross presentation prior to sentence presentation, 2) the sentence presentation epochs which were each 200 ms long and included a sentence presentation onset epoch, a sentence presentation end epoch, a sentence presentation offset epoch, and an epoch preceding the Go cue to repeat) (Figure 4 Panel A) (Table 1). A modulation of activity measure (ΔRMS = RMS of μV_(sentence presentation segment)_ – RMS of μV_(fixation segment)_) was computed for each trial. In the Supplementary Material section we describe a similar analysis using the individually normalized difference as the dependent variable; the results reflect similar patterns.

**Figure 2.**
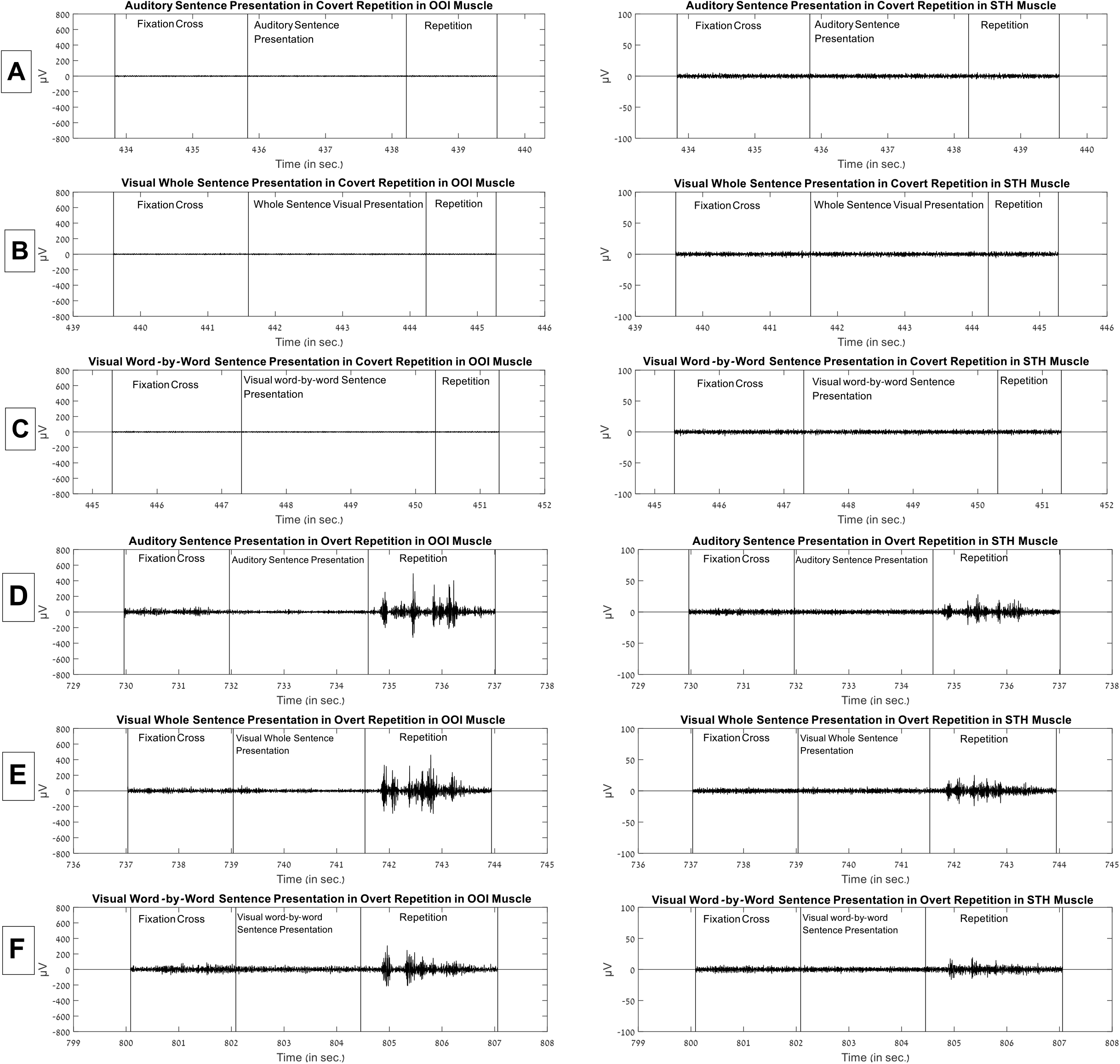
sEMG activity separately for the covert and overt conditions of the task. OOI – Orbicularis Oris Inferior, left panels; STH - Sternohyoid, right panels. Each panel presents a single trial. A,B,C: auditory, visual whole-sentence, and visual word-by-word presentation, respectively, for covert repetition; D, E, F: auditory, visual whole-sentence, and visual word-by-word presentation, for overt repetition. The vertical lines indicate (from left-to-right) the fixation onset, sentence presentation onset, the GO cue onset, repetition completion.

**Figure 3:**
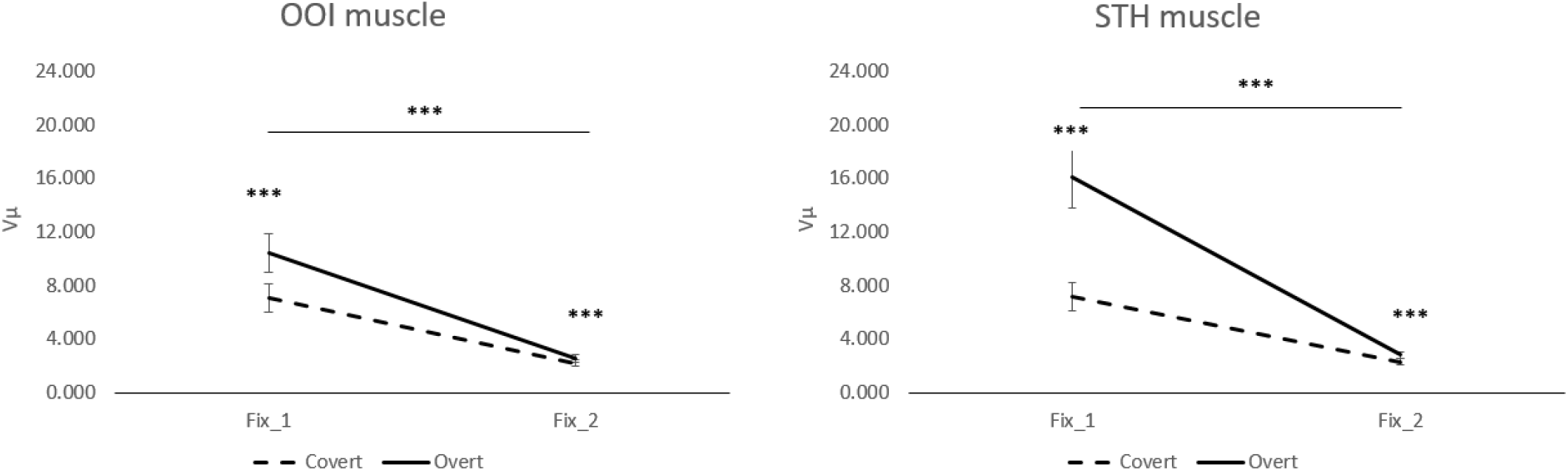
Group average sEMG activity (RMS) at 2 intervals during the fixation epoch. Data are shown for the 2 intended outputs, Overt (Ov) vs. Covert (Cov) in 2 intervals of the fixation epoch (Fix_1 – initial 200 ms; Fix_2 – final 200ms). In the OOI muscle (left panel), there was a significant main effect of Intended Output (F_(1,19)_=14.634, p=.001, η_p_^2^= .435) due to stronger activation in the overt condition (mean diff. = 1.856, p=.001). There was also a significant main effect of Fixation interval (F_(1,19)_=32.615, p=.000, η_p_^2^= .632) due to more activation in the 1^st^ fixation interval compared to the 2^nd^ fixation interval (mean diff. = 6.349, p=.000). In addition, there was a significant interaction of Intended Output and Fixation interval (F_(1,19)_=6.709, p=.018, η_p_ ^2=^ .261). The interaction reflected a smaller difference between the 1^st^ and 2^nd^ fixation intervals in the Covert condition (F_(1,19)_=22.621, p=.000, η_p_^2^=.544) compared to a larger effect in the Overt condition (F_(1,19)_=29.277.14, p=.000, η_p_^2^=.606). Similarly, in the STH muscle (right panel) there was a main effect of Intended Output (F_(1,19)_=24.456, p=.000, η_p_^2^=.563), with more activation prior to overt speech (mean diff.=4.692, p=.000). There was also a main effect of Fixation interval (F_(1,19)_=36.746, p=.000, η_p_^2^=.659), with more activation in the 1^st^ fixation interval compared to the 2^nd^ fixation interval (mean difference=9.059, p=.000). Additionally, there was a significant interaction of Intended Output and Fixation interval (F_(1,19)_=17.326, p=.001, η_p_^2^= .477). The interaction reflected a smaller difference between the 1^st^ and 2^nd^ fixation intervals in the Covert condition (F_(1,19)_=21.888, p=.000, η_p_^2^=.535) compared to a larger effect in the Overt condition (F_(1,19)_=32.460.14, p=.000, η_p_^2^=.631). *p <0.05, ** p <0.01, *** p <0.00. Error bars - standard errors.

**Figure 4:**
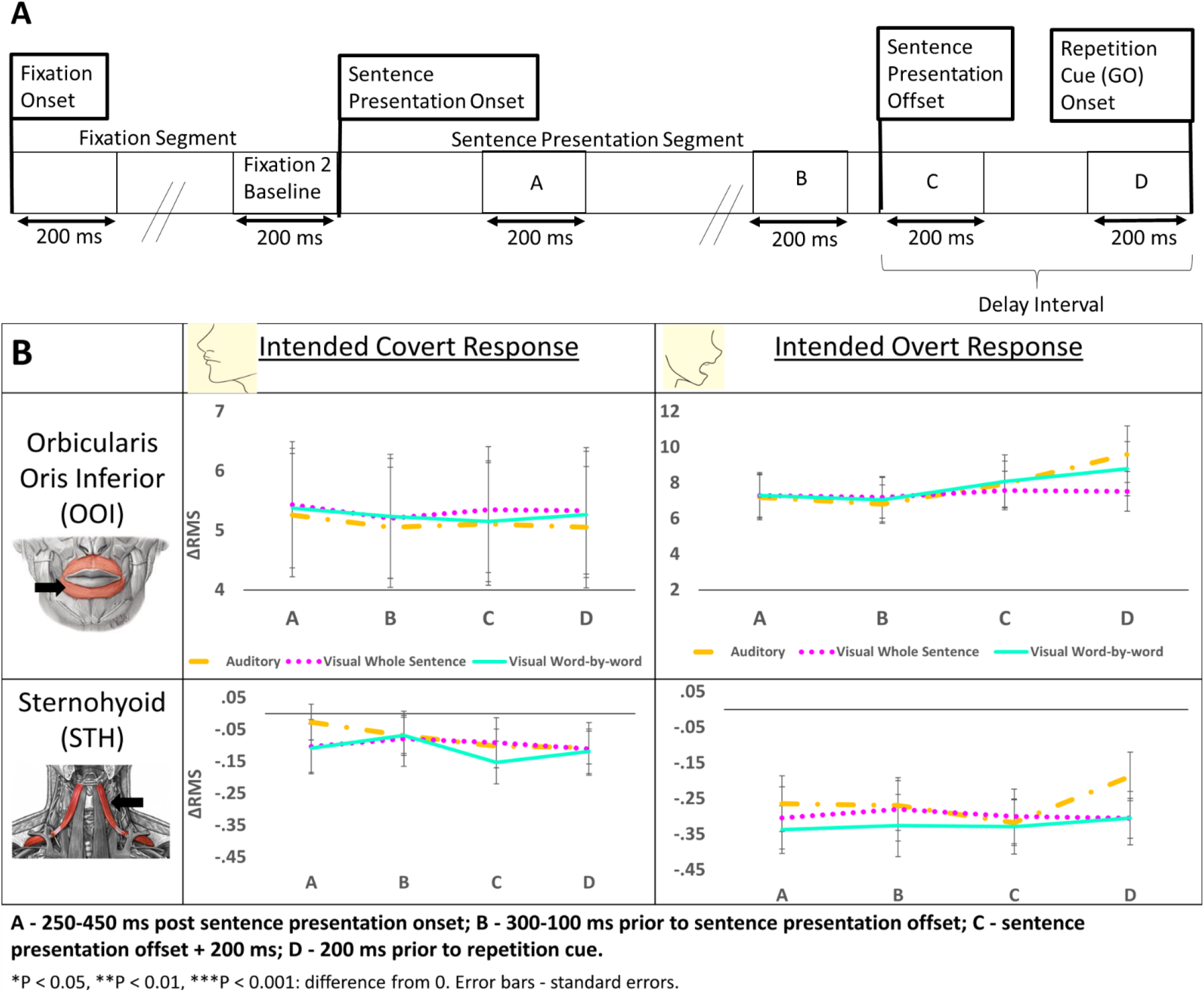
Timeline of the intervals within each trial and ΔRMS Analysis of Variance–. **A)** 4 intervals within the sentence presentation and delay period are shown: intervals A, B, C and D. The delay period includes intervals C and D. **B)** ΔRMS values (vis-a-vis the 2^nd^ fixation interval) of each of the intervals (A, B, C, and D) are shown separately for each of the Intended Output conditions (Covert vs. Overt). * p <0.05, ** p <0.01, *** p< 0.001. Error bars - standard errors.

**Table 1.**
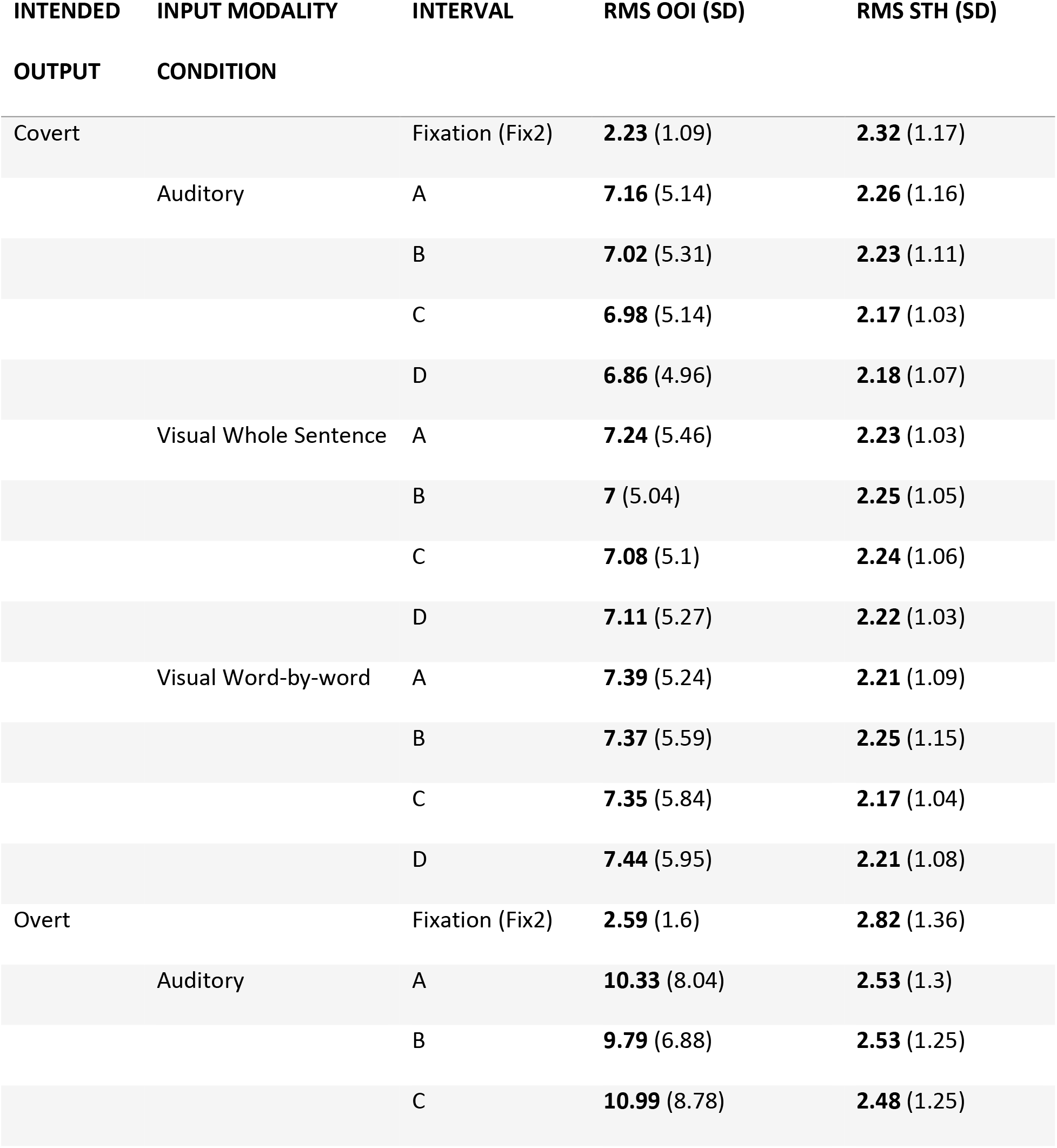
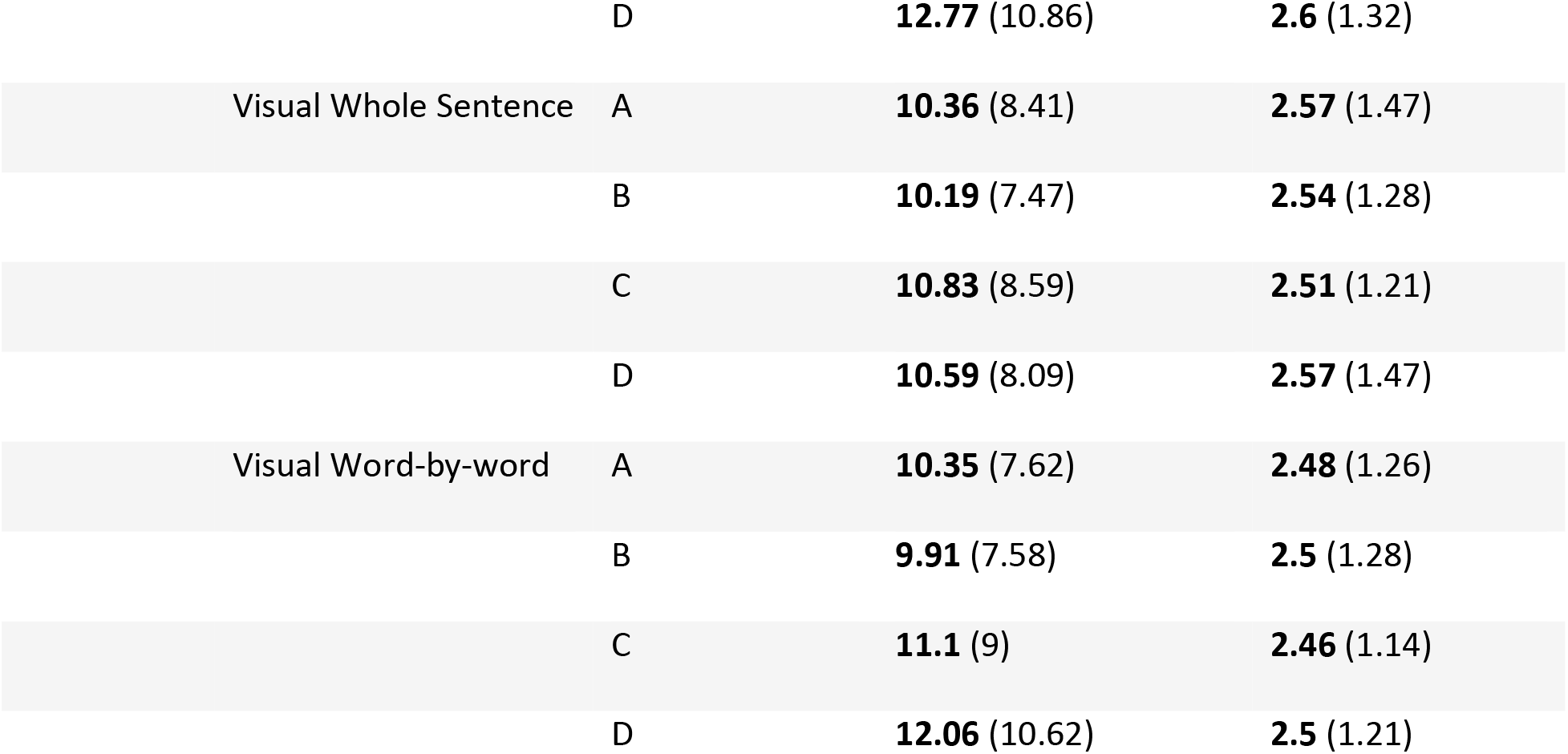
Mean and standard deviation for the root mean square (RMS) of the sEMG activity (μV) within each epoch (fixation and sentence presentation intervals), in each muscle (Orbicularis Oris Inferior - OOI, and Sternohyoid - STH). The RMS values are shown for each of the final 200 ms of the fixation cross segment (Fix2) separately for two experiment blocks (Covert and Overt), and for each of the four intervals in each of the six conditions of the experimental design (A_Cov = Auditory presentation, covert repetition intended; V1_Cov = visual whole-sentence presentation, covert repetition intended; V2_Cov = visual word-by-word sentence presentation, covert repetition intended; A_Ov = Auditory presentation, overt repetition intended; V1_Ov = visual whole-sentence presentation, overt repetition intended; V2_Ov = visual word-by-word sentence presentation, overt repetition intended).

### Fixation Epoch

An analysis of the fixation epoch was undertaken following the findings of an ECoG study, where data was recorded during a single word repetition task (Pei et al., 2016)^28^. Of relevance here, is that in this ECOG study^28^, there was a difference in the signal dynamics during baseline, between the overt and the covert conditions (see their Figure 8, p. 2968). We therefore examined the dynamics within our baseline condition in both muscles. Figure 3 shows the changes in RMS activation values between the first and the last 200 ms of the fixation epoch (which lasted 2 sec). In both muscles, activation tended to decrease across the fixation epoch, and consistently there was higher activity, during fixation, when an overt response was required. Given these dynamics within the fixation epoch, we used the value in the second fixation interval as the baseline in the following analyses.

### Sentence Presentation Epoch Analyses

The analyses of the sentence presentation epoch compared muscle activity (relative to the second interval of the fixation epoch) in four intervals of 200 ms each (Figure 4A), separately for each of the muscles – OOI and STH. These epochs included an interval related to the onset of sentence presentation (A), to the end of sentence presentation (B), an interval beginning at the offset of the sentence presentation (beginning of the temporally jittered delay period) (C), and the interval just before the Go signal (end of the delay period) (D) (Figure 4A). The RMS values of the sEMG activity (μV) were computed for each of these four pre-Go (start repetition) intervals, and a relative activity measure was computed by subtracting the RMS (μV) value of the second fixation epoch.

A 3-way repeated measures ANOVA of Intended Output (Covert vs. Overt), Input Type (A, V1, V2), and Interval (A, B, C and D) in the OOI muscle, which resulted in a significant main effect of Intended Output (F_(1, 19)_= 9.302, p=.007, η_p_^2^=.329), and a significant main effect of Interval (F_(1.825, 34.680)_= 7.288, p=.003, η_p_^2^=.277) (Huynh-Feldt corrected). The main effect of Intended Output was the result of stronger activation in the OOI muscle when overt responses were intended compared to intended covert responses (mean diff.: 3.137; p = .007). The main effect of Interval resulted from Interval D having significantly stronger activation compared to Interval B (mean diff.: .947; p = .007), and marginally significantly stronger activation compared to Interval C (mean diff.: .424; p = .076). Interval C also had significantly stronger activation compared to Interval B (mean diff.: .524; p = .019). In addition, there was a significant Intended Output X Interval interaction (F_(2.169, 41.209)_= 8.900, p=.001, η_p_^2^=.319) as well as an Intended Output X Input Type X and Interval interaction (F_(3.472, 65.959)_= 3.019, p=.029, η_p_^2^=.137) (Huynh-Feldt corrected).

To examine the differential effects of modality, we tested the interaction of Input Type with Interval separately for the two conditions of Intended Output. In the Covert condition, there was a significant main effect of Input Type (F_(1.810, 34.390)_= 3.722, p=.038, η_p_^2^=.134, Huynh-Feldt corrected) so that Auditory presentation had tended to result in lower activation compared to both visual presentation conditions. In the Overt condition, there was a significant main effect of Interval (F_(1.959, 37.230)_= 8.314, p=.001, η_p_^2^=.304, Huynh-Feldt corrected), as well as a significant Input Type X Interval interaction (F_(3.798, 72.170)_= 2.554, p=.049, η_p_^2^=.118, Huynh-Feldt corrected). The significant main effect of Interval was due to significantly stronger activation in Interval D compared to Interval B (mean diff.: 1.874; p = .005), and marginally significantly stronger activation of Interval D compared to both Interval A (mean diff.: 1.480; p = .078) and Interval C (mean diff.: .838; p = .065). In addition, there was significantly more activation in Interval C compared to Interval B (mean diff.: 1.036; p = .010). A separate 1-way ANOVA was conducted for each of the input types (Auditory, Visual whole sentence, and Visual Word-by-word) to look into the source of the Input Type X Interval interaction. In the Auditory input condition, there was a significant main effect of Interval (F_(1.787, 33.957)_= 7.945, p=.002, η_p_^2^=.295, Huynh-Feldt corrected) due to significantly stronger activation in Interval D compared to Interval B (mean diff.: 3.060; p = .020), marginally significantly stronger activation compared to both Interval A (mean diff.: 2.510; p = .055) and compared to Interval C (mean diff.: 1.857; p = .090). In the Visual whole sentence input condition, there was no significant main effect of Interval, and in the Visual Word-by-word condition there was a significant main effect of Interval (F_(2.156, 40.969)_= 5.197, p=.008, η_p_^2^=.215, Huynh-Feldt corrected) which was due to significantly stronger activation in Interval D compared to Interval B (mean diff.: 2.023; p =.030).

The ΔRMS values of the muscle activity during all intervals indicated that the STH muscle was inhibited throughout the sentence presentation epoch (see Table 1). A 3-way repeated measures ANOVA of Intended Output (Covert vs. Overt), Input Type (A, V1, V2), and Interval (A, B, C and D) in the STH muscle was computed. There was a marginally significant main effect of Intended Output (F_(1, 19)_= 3.699, P=.070, η_p_^2^=.163) due to marginally significantly stronger inhibition in overt intended responses compared to covert intended responses (mean diff.: .196; P= .070).

One-sample t-tests were run on all conditions of the design (Intended Output: Covert and Overt(2) X Input Type: Auditory, Visual whole sentence, and Visual word-by-word (3) X Interval (A,B, C, D) (4)), separately for each of the muscles (OOI and STH). All conditions, except for Interval A of auditory sentence presentation in the Covert intended output condition in the STH, were significantly different from baseline. In the OOI, all values were positive (i.e. activation), and in the STH all values were negative (i.e. inhibition).

## 3. Discussion

The results of the current study show that differential inhibition as a function of whether to subsequently vocalize or not, occurs in the STH, with more inhibition when overt responses were intended. In addition, the OOI appears to be more activated compared to baseline, and again, the effect was more pronounced when an overt response was intended. The modality of presentation interacted with both intended output and with the interval: Articulatory muscle activation in the OOI in the covert condition revealed overall less activation during sentence presentation in the auditory than the visual condition. When overt repetition was required, activation was significantly stronger in the last interval (prior to Go cue to repeat) for both auditory and visual word-by-word sentence presentation, but not for the visual whole sentence presentation condition.

One of the important findings of the current study is the dissociation between pattern of activation in the OOI and the STH. The articulatory motor system is often viewed as a singular, synched mechanism. Especially in studies that examine neural activation during speech production. However, these results reflect the complexity of the articulatory system in that facial musculature involved in articulation is activated prior to repetition (in comparison to baseline) while laryngeal musculature involved in articulation is inhibited prior to repetition (in comparison to baseline).

These findings suggest that motor activation of facial articulatory musculature occurs during speech perception, and they align with findings from studies of the neural mechanisms of speech which also show this. For example, Pulvermuller et al. (2006)^29^ found that areas of the precentral gyrus associated with tongue and lip movements during phoneme production were also activated in a somatotopic manner during perception of the same syllables (i.e. when subjects listened to the same phonemes). Feedback and control models of speech production, such as the HSFC model and the DIVA model also posit the involvement of motor cortical areas (namely, the inferior frontal gyrus) during speech perception(Hickok et al., 2011; Guenther & Hickok, 2016)^10,11^.

Additionally, these results show that activation was affected by modality of presentation during the delay period, after sentence presentation and before the Go cue to initiate repetition. Feedback and control models of speech production, such as the HSFC model and the DIVA model, posit that acoustic targets guide articulatory motor production. According to the DIVA model, an auditory target map is formed prior to speech production initiation guiding forward predictions as to how the production should sound^11^. The auditory target map can be manipulated by presenting variations of the acoustic targets. For example, in Tourville et al. (2008)^30^ perturbed speech was presented to subjects as their own produced speech in several, random trials, and the result was a compensation in production of the syllables in the direction opposite of the perturbation. Thus, according to the model, differences in auditory stimuli to be repeated (i.e. auditory vs. visually presented sentences) can result in different motor output. When stimuli are presented visually, the auditory representation of the linguistic content is generated internally. This is the difference between the auditory (external auditory representation) and visually (internal auditory representation) presented sentences. The HFSC model assumes internal auditory representation as auditory targets required for repetition (i.e. visually presented sentences to be repeated), production will be different compared to when repetition of an external auditory target is required (Hickok et al., 2011)^10^. The effect of input type presented here, supports the assumptions of both feedback models.

The effect of input type replicate results of EcoG studies. For example, the study by Pei et al. (2011)^28^, found that frontal activity patterns were significantly different during the input stage of a single-word repetition task, as a functions of modality of presentation (see their figure 8 p.2968).

The finding that the intended response (Covert vs. Overt) affected muscle activation patterns in the stimulus presentation epoch and during the delay intervals, are similar to those reported by Cogan et al. (2017)^31^ in a ECoG study of verbal Working Memory (vWM). These results support the notion that processing phonological input for repetition or production purposes is active (rather than passive) (Cogan et al., 2017; Ojemann et al. 2009; Zamora et al., 2016)^31, 32, 33^.

The results of the current study present similar articulatory muscle activation trends for covert and overt speech. In the OOI, both covert and overt intended responses elicited activation throughout input perception and the delay intervals. In the STH, both covert and overt intended responses elicited inhibition throughout input perception and the delay intervals. In both cases, when overt responses were intended, activity in the OOI or inhibition in the STH was stronger compared to when covert responses were intended. This result can be linked to the findings of Brumberg et al. (2016)^34^ that reported ECoG recordings in participants instructed to read a familiar text aloud or silently. They found left fronto-motor activity at 440-240 ms prior to the initiation of production, in both the overt and covert speech production conditions. However, this activity was significantly higher when overt speech production was intended compared to the covert speech production condition. The findings of the current study raise the possibility that the left fronto-motor areas dynamics may relate, directly or indirectly, to the differential dynamics of inhibition of the articulatory musculature in the language perception phase. However, our measurements were purely motor; we have no way of knowing their origins in the central nervous system.

The current results may be construed as reflecting a type of ‘preparatory inhibition’ in relation to elicited speech (Conrad et al., 1983; Richartz et al., 2010; Kolářová et al., 2016; Duque et al., 2017; Lebon et al., 2016) ^23–25,5–36^. However, there is a difference in the time scales. For example, Lebon et al. (2016)^36^, found inhibitory preparatory processes only 200 ms (but not 500 ms or 900 ms) prior to an imperative signal to move either the left or right index fingers in a delayed response task. Unlike the trend for increased inhibition closer to the Go signal, found by Lebon et al. (2016)^36^, in the current study, inhibition tended to be consistent from the onset of sentence presentation up to the final 200 ms interval before the GO signal. A positive relationship between the magnitude of pre-movement depression of tonic muscle activity and the subsequent phasic innervation burst has been suggested for ballistic arm movements (Conrad et al., 1983)^23^. Our findings may be linked to the findings of Conrad et al. (1983)^23^, if one considers the sentence presentation (input) epoch in the current study as a predictive phase necessarily preceding the actual Go signal. With the caveat of a different time scale and resolution, the current results show that the intended speech action (overt or covert) was a major factor in determining the degree of inhibition or activation. However, the extended time course of muscle modulation and the fact that it occurred from the very beginning of the sentence presentation epoch, when both a covert and overt response was intended, suggest a pattern of modulation that differs from the pattern of inhibition described in non-language manual responses.

The notion that inhibition in the STH muscle can be linked to motor preparartion of speech production is supported by findings of neuroimaging studies of the LMC. A fMRI study by Symonian et.al. (2009)^17^ used a syllable repetition task and a controlled breathing task to examine the involvement of the LMC in controlled breathing and in speech. They found bilateral activation of the LMC during the controlled breathing task, and a left lateralized activation of the LMC during the syllable repetition task. The authors also observed functional coupling between the LMC and Inferior Frontal Gyrus (IFG), and together with the finding that evoked activity of the LMC during IFG electrical stimulation (Greenlee et al., 2004)^37^, they concluded that the processing of any speech production component requires a functional link between these two brain regions to enable speech motor preparation. The finding that STH muscle activation is modulated by the intention to speak or not, supports this hypothesis.

## Conclusions

In a more general context, the current findings are also consistent with previous findings showing that planned, intended, motor actions can significantly affect ongoing actions; e.g., co-articulation, the finding that skilled speakers generate the initial phonemes of a sequence differentially, depending on the final phonemes (Kühnert & Nolan, 1999) ^38^. This has been has shown in a non linguistic task as well. Rozanov et al. (2010)^39^ showed that the performance of the initial movements of a well-trained sequence of finger movements was compromised by the intention to subsequently omit a movement. In the current study, anticipatory effects were evident at the stage of input, before articulation actually occurred.

The current results indicate that, prior to repetition, the articulatory system was differentially activated in the different muscle systems during speech perception and reading. Muscle activity was modulated by the intention to subsequently vocalize the input or not. Thus, the intention to act in the future (in terms of voicing) was a significant factor in articulatory muscle activity during language perception. The data of the current study suggest that the role of intention in language production during reading and listening, is dynamic, continuous, and contextual.

## References

1. Liberman, A. M. Some results of research on speech perception. The Journal of the Acoustical Society of America, 29(1), 117–123 (1957).

2. Liberman, A. M., Cooper, F. S., Shankweiler, D. P., & Studdert-Kennedy, M. Perception of the speech code. Psychological review, 74(6), 431 (1967).

3. Wilson, S. M., Saygin, A. P., Sereno, M. I., & Iacoboni, M. Listening to speech activates motor areas involved in speech production. Nature neuroscience, 7(7), 701–702 (2004).

4. Pulvermüller, F., & Fadiga, L. Active perception: sensorimotor circuits as a cortical basis for language. Nature Reviews Neuroscience, 11(5), 351–360 (2010).

5. Yuen, I., Davis, M. H., Brysbaert, M., & Rastle, K. Activation of articulatory information in speech perception. Proceedings of the National Academy of Sciences, 107(2), 592–597 (2010).

6. Skipper, J. I., Devlin, J. T., & Lametti, D. R. The hearing ear is always found close to the speaking tongue: review of the role of the motor system in speech perception. Brain and language, 164, 77–105 (2017).

7. Hickok, G., & Poeppel, D. The cortical organization of speech processing. Nature Reviews Neuroscience, 8(5), 393–402 (2007).

8. Poeppel, D., & Hickok, G. Towards a new functional anatomy of language. Cognition, 92(1), 1–12 (2004).

9. Scott, S. K., McGettigan, C., & Eisner, F. A little more conversation, a little less action— candidate roles for the motor cortex in speech perception. Nature Reviews Neuroscience, 10(4), 295–302 (2009).

10. Hickok, G., Houde, J., & Rong, F. Sensorimotor integration in speech processing: Computational basis and neural organization. Neuron, 69(3)., 407–422 (2011).

11. Guenther, F. H., & Hickok, G. (2016). Neural models of motor speech control. In Neurobiology of language (pp. 725–740). Academic Press.

12. Galantucci, B., Fowler, C. A., & Turvey, M. T. The motor theory of speech perception reviewed. Psychonomic bulletin & review, 13(3), 361–377 (2006).

13. Glenberg, A. M., & Gallese, V. Action-based language: A theory of language acquisition, comprehension, and production. Cortex, 48(7), 905–922 (2012).

14. Logan, G. D. Automaticity and reading: Perspectives from the instance theory of automatization. Reading & Writing Quarterly: Overcoming Learning Difficulties, 13(2), 123–146 (1997).

15. Augustinova, M., & Ferrand, L. Automaticity of word reading: Evidence from the semantic Stroop paradigm. Current Directions in Psychological Science, 23(5), 343–348 (2014).

16. Jürgens, U. (2002). Neural pathways underlying vocal control, in «Neuroscience & Biobehavioral Reviews», 26.

17. Simonyan, K., Ostuni, J., Ludlow, C. L., & Horwitz, B. (2009). Functional but not structural networks of the human laryngeal motor cortex show left hemispheric lateralization during syllable but not breathing production. Journal of Neuroscience, 29(47), 14912–14923.

18. Simonyan, K. (2014). The laryngeal motor cortex: its organization and connectivity. Current opinion in neurobiology, 28, 15–21.

19. Jacobson, E. Electrical measurements of neuromuscular states during mental activities. American Journal of Physiology--Legacy Content, 97(1), 200–209 (1931).

20. Livesay, J., Liebke, A., Samaras, M., & Stanley, A. Covert speech behavior during a silent language recitation task. Perceptual and motor skills, 83(3 suppl), 1355–1362 (1996).

21. McGuigan, F. J., & Dollins, A. B Patterns of covert speech behavior and phonetic coding. The Pavlovian journal of biological science, 24(1), 19–26 (1989).

22. Rapin, L., Dohen, M., Polosan, M., Perrier, P., & Lœvenbruck, H. An EMG study of the lip muscles during covert auditory verbal hallucinations in schizophrenia. Journal of Speech, Language, and Hearing Research, 56(6), 1882–1893 (2013).

23. Conrad, B., Benecke, R., & Goehmann, M. Premovement silent period in fast movement initiation. Experimental Brain Research, 51(2), 310–313 (1983).

24. Richartz, C., Lévénez, M., Boucart, J., & Duchateau, J. Initial conditions influence the characteristics of ballistic contractions in the ankle dorsiflexors. European journal of applied physiology, 110(4), 805–814 (2010).

25. Kolářová, B., Krobo, A., Kolář, P., Hluštík, P., & Polehlová, K. Muscle Activity Inhibition in Lower Limb Muscles during Gait Imagery Tasks. Current Research in Motor Control V, 123 (2016).

26. Whiting, C., Shtyrov, Y., & Marslen-Wilson, W. Real-time functional architecture of visual word recognition. Journal of Cognitive Neuroscience, 27(2), 246–265 (2014).

27. Stepp, C. E. Surface electromyography for speech and swallowing systems: Measurement, analysis, and interpretation. Journal of Speech, Language, and Hearing Research: JSLHR, 55(4)., 1232–1246 (2012).

28. Pei, X., Leuthardt, E. C., Gaona, C. M., Brunner, P., Wolpaw, J. R., & Schalk, G. Spatiotemporal dynamics of electrocorticographic high gamma activity during overt and covert word repetition. Neuroimage, 54(4), 2960–2972 (2011).

29. Pulvermüller, F., Huss, M., Kherif, F., del Prado Martin, F. M., Hauk, O., & Shtyrov, Y. (2006). Motor cortex maps articulatory features of speech sounds. Proceedings of the National Academy of Sciences, 103(20)., 7865–7870.

30. Tourville, J. A., Reilly, K. J., & Guenther, F. H. (2008). Neural mechanisms underlying auditory feedback control of speech. Neuroimage, 39(3)., 1429–1443.

31. Cogan, G. B., Iyer, A., Melloni, L., Thesen, T., Friedman, D., Doyle, W.,…& Pesaran, B. (2017). Manipulating stored phonological input during verbal working memory. Nature neuroscience, 20(2)., 279–286.

32. Ojemann, G. A., Schoenfield-McNeill, J., & Corina, D. (2009). The roles of human lateral temporal cortical neuronal activity in recent verbal memory encoding. Cerebral Cortex, 19(1)., 197–205.

33. Zamora, L., Corina, D., & Ojemann, G. (2016). Human temporal cortical single neuron activity during working memory maintenance. Neuropsychologia, 86, 1–12.

34. Brumberg Jonathan S., et al. “Spatio-Temporal Progression of Cortical Activity Related to Continuous Overt and Covert Speech Production in a Reading Task.” PloS one, 11.11 (2016).

35. Duque, J., Greenhouse, I., Labruna, L., & Ivry, R. B. Physiological markers of motor inhibition during human behavior. Trends in neurosciences, 40(4), 219–236 (2017).

36. Lebon, F., Greenhouse, I., Labruna, L., Vanderschelden, B., Papaxanthis, C., & Ivry, R. B. Influence of delay period duration on inhibitory processes for response preparation. Cerebral Cortex, 26(6), 2461–2470 (2015).

37. Greenlee, J. D., Oya, H., Kawasaki, H., Volkov, I. O., Kaufman, O. P., Kovach, C.,…& Brugge, J. F. (2004). A functional connection between inferior frontal gyrus and orofacial motor cortex in human. Journal of neurophysiology, 92(2)., 1153–1164.

38. Kühnert, B., & Nolan, F. The origin of coarticulation. Coarticulation: Theory, Data and Techniques (1999).: 7–30.

39. Rozanov, S., Keren, O., & Karni, A. The specificity of memory for a highly trained finger movement sequence: Change the ending, change all. Brain Research, 1331, 80–87 (2010).

