## Supplementary Material for "Modulation of articulation muscles during listening and reading: a matter of intention to speak out loud"

### Sentence Presentation Epoch Analyses – Normalized ΔRMS

As mentioned in the methods section, RMS values of sEMG activity (µV) were computed for each of the intervals, and a relative activity measure was computed by subtracting the RMS (µV) value of Fix2 from each of the RMS(µV) values of the intervals. However, for normalized values we divided the difference values by the RMS (µV) value of Fix2 and multiplied the ratio by 100. Therefore, the normalized value reflected the percent change in the interval compared to the Fix2 interval.

The analysis examined the dynamics of muscle activity (relative to the second interval of the fixation epoch) in four intervals of 200 ms each (Figure 4A), separately for each of the muscles – OOI and STH.

A 3-way repeated measures ANOVA of Intended Output (Covert vs. Overt), Input Type (A, V1, V2), and Interval (A, B, C and D) in the OOI muscle resulted in a significant main effect of Intended Output ($F_{(1, 19)}$= 6.723, p=.018, $\eta_p^2$=.261), and a significant main effect of Interval ($F_{(1.964, 33.567)}$= 6.903, p=.003, $\eta_p^2$=.266) (Huynh-Feldt corrected). The main effect of Intended Output was the result of stronger activation in the OOI muscle when overt responses were intended compared to intended covert responses (mean diff.: 136.941; p = .018). The main effect of Interval resulted from Interval B having significantly more diminished activation compared to Interval D (mean diff.: 45.632; p = .012), and compared to Interval C (mean diff.: 7.598; p = .023). In addition, all three interactions were also significant. There was a significant Intended Output X Interval interaction ($F_{(2.221, 42.190)}$= 8.971, p=.000, $\eta_p^2$=.321), a  significant Intended Output X Input Type ($F_{(4.301, 81.710)}$= 2.541, p=.042, $\eta_p^2$=.118), and an Intended Output X Input Type X and Interval interaction ($F_{(3.494, 66.389)}$= 3.533, p=.015, $\eta_p^2$=.157) (Huynh-Feldt corrected).

To examine the effect of modality, we tested the interaction of Input Type with Interval separately for the two conditions of Intended Output. In the Covert intended response condition, there was a significant main effect of Interval ($F_{(2.220, 42.174)}$= 3.144, p=.048, $\eta_p^2$=.142, Greenhouse-Geisser corrected). Planned contrasts show that this effect is due to a marginally significant difference between Interval A and Interval B, so that there was stronger activation in Interval A compared to Interval B ($_{((1, 19)}$= 3.881, p=.064, $\eta_p^2$=.170). In the Overt condition, there was a significant main effect of Interval ($F_{(2.067, 39.264)}$= 8.068, p=.001, $\eta_p^2$=.298, Huynh-Feldt corrected), as well as a significant Input Type X Interval interaction ($F_{(3.750, 71.252)}$= 3.123, p=.022, $\eta_p^2$=.141, Huynh-Feldt corrected). The significant main effect of Interval was due to significantly stronger activation in Interval D compared to Interval B (mean diff.: 92.470; p = .008), and marginally significantly stronger activation of Interval D compared to both Interval A (mean diff.: 70.956; p = .070). In addition, there was significantly more activation in Interval C compared to Interval B (mean diff.: 51.526; p = .009). A separate 1-way ANOVA was conducted for each of the input types (Auditory, Visual whole sentence, and Visual Word-by-word) to look into the source of the Input Type X Interval interaction. In the Auditory input condition, there was a significant main effect of Interval ($F_{(1.609, 30.567)}$= 8.086, p=.003, $\eta_p^2$=.299, Huynh-Feldt corrected) due to significantly stronger activation in Interval D compared to Interval A (mean diff.: 126.177; p = .046), Interval B (mean diff.: 153.404; p = .029) and Interval C (mean diff.: 103.010; p = .049). In the Visual whole sentence input type condition, there was no significant main effect of Interval, and in the Visual Word-by-word condition there was a significant main effect of Interval ($F_{(1.893, 35.963)}$= 5.493, p=.009, $\eta_p^2$=.224, Huynh-Feldt corrected) which was due to significantly more diminished activation in Interval B compared to Interval C (mean diff.: 62.444; p = .050) and Interval D (mean diff.: 106.293; p = .042).

The ΔRMS values of the muscle activity during all intervals indicated that the STH muscle was inhibited throughout the sentence presentation epoch. A 3-way repeated measures ANOVA of Intended Output (Covert vs. Overt), Input Type (A, V1, V2), and Interval (A, B, C and D) in the STH muscle was computed. There was a marginally significant main effect of Intended Output ($F_{(1, 19)}$= 3.993, p=.060, $\eta_p^2$=.174) due to marginally significantly stronger inhibition in overt intended responses compared to covert intended responses (mean diff.: 5.751; p = .060).
